# Habitat fragmentation and species diversity in competitive species communities

**DOI:** 10.1101/363234

**Authors:** Joel Rybicki, Nerea Abrego, Otso Ovaskainen

## Abstract

Habitat loss is one of the key drivers of the ongoing decline of biodiversity. However, ecologists still argue about how fragmentation of habitat (independent of habitat loss) affects species richness. The recently proposed habitat amount hypothesis posits that species richness only depends on the total amount of habitat in a local landscape. On the other hand, different empirical studies report contrasting patterns: some find positive and others negative effects of fragmentation *per se* on species richness.

To explain this apparent disparity, we devise a stochastic, spatially-explicit model of competitive species communities in heterogeneous habitats. The model shows that habitat loss and fragmentation have a non-monotone and non-linear effect on the species diversity in competitive communities. When the total amount of habitat is large, fragmentation *per se* tends to increase species diversity, but if the total amount of habitat is small, the situation is reversed: fragmentation *per se* decreases species diversity.

## 1 Introduction

Degradation and loss of natural habitat due to anthropogenic modification and climate change is a key factor contributing to the ongoing sixth extinction event (Tilman et al., 1994; Fahrig, 2003; Thomas et al., 2004; Kuussaari et al., 2009; Pereira et al., 2010; Butchart et al., 2010; Pimm et al., 2014). Habitat *loss* (reduction of area with suitable habitat) goes hand in hand with habitat *fragmentation* (division of the habitat into several parts), where the former is a process causing the latter landscape pattern (Fahrig, 2003; Ewers and Didham, 2005; Wilson et al., 2016). While the decline of biodiversity due to habitat loss is uncontested, the effect of habitat fragmentation *per se* on species richness has been much debated over the past decades (Fahrig, 2003; Ewers and Didham, 2005; Didham et al., 2012; Fahrig, 2013; Hanski, 2015; Haddad et al., 2015; Fahrig, 2017): given the same total amount of habitat, how does the spatial configuration of the habitat, that is, the locations, shapes, and sizes of habitat fragments, influence biodiversity?

Indeed, one of the key questions in conservation biology is whether protecting biodiversity is better achieved using a single large or several small (SLOSS) reserves (Diamond, 1975; Ewers and Didham, 2005). For species that follow classical metapopulation dynamics (Hanski, 1999), the effects of fragmentation and spatial configuration are well-understood from the theoretical perspective (Bascompte and Solé, 1996; Hanski and Ovaskainen, 2000; Ovaskainen, 2002; Hanski and Ovaskainen, 2003; Gilarranz and Bascompte, 2012; Grilli et al., 2015): single species metapopulation theory predicts that increasing fragmentation is detrimental for species – although the response is not necessarily monotone (Ovaskainen, 2002) – assuming that no evolutionary responses take place; see Legrand et al. (2017) for a review of ecoevolutionary responses to habitat fragmentation. On the other hand, increasing connectivity in fragmented landscapes can increase synchrony in metapopulations, and consequently, lead to increased extinction risk (Kahilainen et al., 2018).

However, many species do not necessary follow metapopulation dynamics or the scale at which they do is limited. Moreover, the situation becomes much more muddled when considering species communities that comprise several interacting species. While increasing fragmentation is known to have largely negative effects for metapopulations, this does not necessarily hold for metacommunities of several species. Indeed, while some species (e.g. habitat specialists) may suffer from fragmentation, others may benefit from it (e.g. generalists and edge species) (Henle et al., 2004). Various theoretical models of multispecies communities and empirical evidence suggests that different spatial configurations of the habitat can have both negative and positive effects on species richness (Tilman et al., 1997; Rybicki and Hanski, 2013; Hanski et al., 2013; Hanski, 2015; Haddad et al., 2015; Fahrig, 2017; Thompson et al., 2017; Haddad et al., 2017) depending on the species’ traits together with properties and structure of the landscape (e.g. degree of spatial autocorrelation in habitat types). Nevertheless, current theory still suggests that fragmentation (per se) tends to increase extinctions in predator-prey metapopulations and competitive metacommunities (Tilman et al., 1997).

In a recent meta-analysis of empirical fragmentation studies, Fahrig (2017) concluded that the most significant ecological responses to habitat fragmentation were positive instead of negative. The positive effects of fragmentation have been attributed to numerous causes including – but not limited to – increase in functional connectivity, diversity of habitat types, persistence of predator-prey systems and decrease in intra- and interspecific competition.

In contrast, some studies have reported negative effects including increased risk of extinction due to reduced genetic diversity, environmental and demographic stochasticity in small patches (Ewers and Didham, 2005). Various edge effects have been proposed to have both positive and negative effects (Ewers and Didham, 2005; Fahrig, 2017) depending on the species traits. Furthermore, fragmentation may alter species interactions and community composition, as invasive or pest species may replace the original species pool, increase the transmission and prevalence of disease in small fragments, and the effects of fragmentation can be confounded by the associated time lags (Ewers and Didham, 2005; Haddad et al., 2015). Thus, the effects of fragmentation on species communities remain far from well-understood.

### 1.1 The habitat amount hypothesis

Recently, Fahrig (2013) has proposed the *habitat amount hypothesis*, which postulates that the species richness is best explained by the sample area effect: large areas of habitat tend to support more individuals, and hence, more species (Rosenzweig, 1995). The hypothesis posits that what truly matters is the *total amount* of habitat in a local landscape independent of its spatial configuration at the “appropriate spatial extent of the local landscape” (Fahrig, 2013). Namely, the hypothesis makes the following predictions:

- **Prediction 1:** Given equal-sized sample sites, species richness increases with total amount of habitat in the “local landscape” of the sample site.
- **Prediction 2:** Species richness is independent of the particular patch in which the sample site is located (except with contribution of the patch area to the total area at local landscape).

Fahrig (2013, 2015) has called for a research program to test the hypothesis, and subsequently, the hypothesis has received considerable attention. For example, Haddad et al. (2017) reported that empirical evidence from fragmentation experiments on plant and micro-arthropod communities does not support the hypothesis, but in contrast Arnillas et al. (2017) found that fragmentation may indeed have positive effect on species richness – at least on the short-term – and De Camargo et al. (2018) report that for birds there is no detectable response to fragmentation at the landscape level, but do not rule out the possibility that fragmentation matters at smaller scalers.

In general, it still remains unclear which species and landscape attributes lead to different responses to habitat fragmentation (Fahrig, 2017). Indeed, the habitat amount hypothesis has been strongly criticised due to lack of an underlying mechanistic explanation, which would predict what is the appropriate spatial extent of the local landscape and how species interactions and community structure affects the species richness (Hanski, 2015). Simulation studies have suggested that the appropriate landscape scale for the hypothesis is an order of magnitude (e.g. 4–9 times) higher than the dispersal scale of species (Jackson and Fahrig, 2012; Fahrig, 2013).

In this work, we set out to make better understanding of the effects of fragmentation in competitive communities. In particular, we investigate the habitat amount hypothesis under different landscape and community properties using a spatially-explicit, individual-based community model. Using our model, we assess Predictions (1)–(2) using the tests proposed by Fahrig (2013).

### 1.2 Prior work on metacommunity models

We focus on species dynamics at the landscape level in fragmented environments, that is, metacommunity dynamics. While a metapopulation is a collection of populations (of a single species) that reside in discrete patches connected by dispersal, the metacommunity concept is its multispecies analogue (Hanski, 1999): patches are inhabited by several (possibly interacting) species and the metacommunity dynamics are governed by various coexistence processes (Leibold et al., 2004). Typically, theoretical studies on metacommunities have generalised patch-based metapopulation models into multispecies models with discrete patches in both spatially-implicit (Tilman et al., 1994; Loreau et al., 2003; Mouquet and Loreau, 2003; Wang and Loreau, 2016) and spatially-explicit models (Rybicki and Hanski, 2013; Matias et al., 2014; Thompson et al., 2014; Fournier et al., 2016; Thompson et al., 2017). The interspecies interactions are typically competitive (e.g. competition on space or a shared limiting resource) (Tilman et al., 1994; Loreau et al., 2003; Mouquet and Loreau, 2003; Wang and Loreau, 2016; Matias et al., 2014; Thompson et al., 2014; Fournier et al., 2016; Thompson et al., 2017), but also mutualistic (e.g. plant-pollinator systems) (Klausmeier, 2001; Prakash and de Roos, 2004; Fortuna and Bascompte, 2006) and trophic interactions (Pillai et al., 2010) have been considered.

A relatively large proportion of the theoretical work on metacommunities has focused on understanding how e.g. coexistence mechanisms and dispersal maintain species diversity and stability (Lehman and Tilman, 2000; Loreau et al., 2003; Mouquet and Loreau, 2003; Gravel et al., 2006; Logue et al., 2011; Haegeman and Loreau, 2014; Wang and Loreau, 2016; Gravel et al., 2016). While less common, models have also been used to investigate how landscape structure, habitat loss and fragmentation influence species richness in metacommunities (Tilman et al., 1997; Prakash and de Roos, 2004; Rybicki and Hanski, 2013; Matias et al., 2014; Thompson et al., 2014, 2017; Xu et al., 2018).

As usual, these models make various assumptions for the sake of tractability: some completely ignore species interactions and rely on species-sorting mechanisms (Rybicki and Hanski, 2013), limit to only pairs of interacting species (Klausmeier, 2001), ignore spatial heterogeneity, assume patch-based metapopulation dynamics (Tilman et al., 1997; Prakash and de Roos, 2004; Rybicki and Hanski, 2013; Matias et al., 2014) or lack demographic and/or environmental stochasticity by employing deterministic, continuous-valued dynamics (Thompson et al., 2014).

### 1.3 Contributions of this work

Our main contributions are summarised as follows:

- **We develop a individual-based, spatially-explicit model of competitive species communities in spatiotemporally varying landscapes.** Our model is mechanistic and relaxes many assumptions made in prior models. For example, we do *not* assume that the species follow metapopulation dynamics or that the habitat consists of discrete patches. Instead, we keep track of individuals who follow stochastic birth-death dynamics in a continuous landscape. Intra- and interspecific competition is achieved via resource-consumer dynamics. The model is extensible and the source code is freely available online^∗^ thus, it can be used to investigate also other spatial aspects of species interactions in community dynamics.
- **We investigate the habitat amount hypothesis following the test criteria laid out by Fahrig (2013).** Using our model, we examine how habitat fragmentation *per se* influences the species richness in competitive species communities. Our model indicates that fragmentation can have *positive* effects on species diversity of competitive habitat specialists if the total amount of habitat in a landscape is large. However, when the total amount of habitat is small, fragmentation yields *negative* effects. Furthermore, we observe that fragmentation has the same *qualitative* effect on both species that are sessile after dispersal and non-sessile species that actively navigate in the landscape in order to find suitable habitat.
- **We show that different analyses of fragmentation effects may lead to apparently contradictory results.** For example, depending on the quality of matrix, species-area relationships may show positive effects at the landscape level, while SLOSS-type analysis indicates negative effects. This suggests that caution is needed when interpreting whether data shows that fragmentation has positive or negative effects on species richness of habitat specialists.

## 2 Model and methods

### 2.1 Spatially-explicit community model

We devise an individual-based community model in continuous time and continuous two-dimensional, spatiotemporally heterogeneous landscapes, where the growth of all species is governed by one or more limiting resources. Our model contains three basic entity types: resource patches, resource particles, and individuals of (each) species. The resource patches produce resource particles into their neighbourhood, and the individuals of the species consume these resource particles. Thus, we obtain intra- and interspecific resource competition yielding density-dependent population growth. The species convert the resources into offspring and all individuals follow birth-death dynamics.

We first define the model dynamics in a pristine landscape without habitat loss. After this, we discuss ways how to include habitat loss and fragmentation into our model. For brevity, we describe the model in the context of a single species consuming a single resource type, but the model is straightforward to generalise to multiple species and resource types.

#### Model structure

Formally, our model is a spatiotemporal point process, or in the mathematical terminology, a Markov evolution in the space of locally finite configurations (see e.g. Ovaskainen et al., 2014). The state of a model at any given time is given by the spatial location of each individual of each type. The dynamics of the model can be described by listing all events that can take place and the rates at which these events occur. These rates can depend on the current spatial configuration of all individuals (e.g. individuals can only consume resources that are within their proximity). The proximity-dependence is described via kernels, that is, functions that tell at which rate two particles at locations *x* = (*x*_1_, *x*_2_) and *y* = (*y*_1_, *y*_2_) react depending on their Euclidean distance dist(*x*, *y*). A kernel *K*(*x*, *y*) = *r* · *f* (*x*, *y*) is defined in terms of a total rate parameter *r* and a density function *f*, which describes how the total rate *r* is distributed accross the space.

We use two types of kernels. A *tophat* kernel *K* with length scale *ℓ* and total rate *r* is defined as

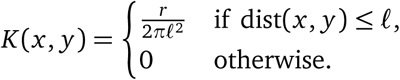

A *Gaussian* kernel *K* with length scale *ℓ* and total rate *r* is given by

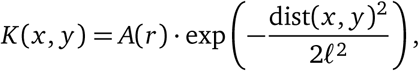

where *A*(*r*) = *r*/(2*πℓ*^2^) is a normalisation constant such that *K* integrates to the value *r*. In the following, we write *ℓ*(*K*) and *r*(*K*) to denote the length scale and total rate of kernel *K* of either type.

#### Patch dynamics

To obtain spatiotemporal variation in resource production rate, we assume that the resource patches follow birth-death dynamics independent of the species. The patch birth-death dynamics are described by the following processes

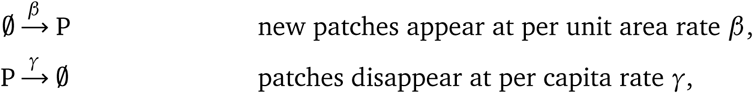

where ∅ denotes “no particle” (i.e. birth of patches does not depend on any entity and the death of a patch simply removes the patch but not the resources it has produced). We assume that the resource patches are circular (but may overlap) and that a resource patch located at *x* produces abiotic resource particles to its surroundings according to the resource generation kernel *G*. Resource particles that are left unconsumed by the species decay and disappear with constant rate. Thus, we have the following processes:

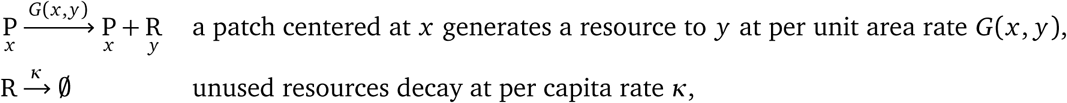

where in the first reaction, the location of a new particle is randomly sampled according to the density function *f* of the kernel *G*(*x*, *y*) = *r* · *f* (*x*, *y*). Note that we can control the spatiotemporal properties of the landscape’s habitat quality by varying the per unit area density *ρ* = *β*/*γ* of habitat patches, the patch turnover rate *γ*, the resource production rate *r* = *r*(*G*), and the radius *ℓ*(*G*) of the patches. First panel of Figure 1 depicts an example snapshot of the resulting habitat structure.

**Figure 1:**
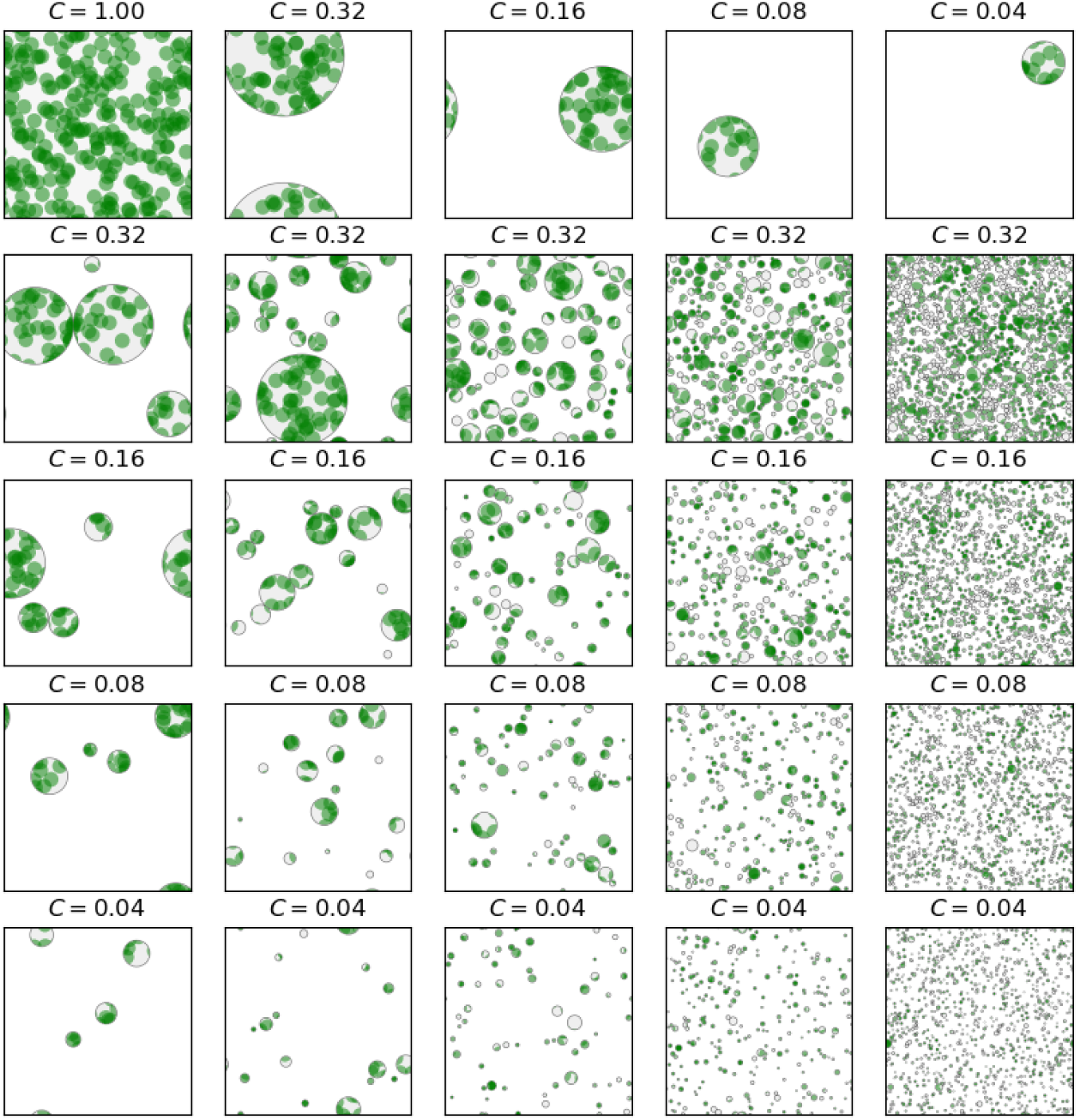
Examples of fragmented landscapes with varying levels of fragmentation and habitat cover. The landscape is a torus (i.e. periodic boundaries). Green cirles are resource patches, which may overlap to create higher resource production rates at some areas (darker green). The filled gray circles represent habitat fragments within which resource patches follow stochastic birth-death dynamics, and within these circles, the gray area represents habitat (currently) without resource patches. The white areas represent the matrix, that is, area which has zero resource production rate.

#### Species dynamics

The species follow birth-death dynamics and individuals have two states. Newborn individuals start out in the *resource*-*deprived* state, and upon consuming a resource particle, they become *resource*-*satiated*; for brevity, we refer to individuals in these states as deprived (D) and satiated (S), respectively. The satiated individuals produce new individuals into their surroundings:

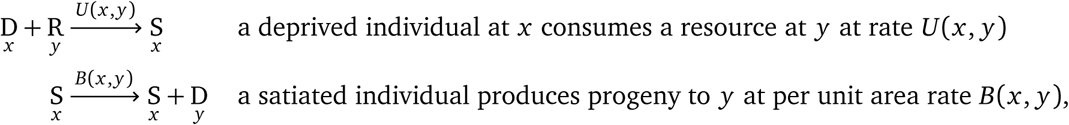

where *U* is a resource utilisation kernel, *B* is a Gaussian birth kernel, and the satiated individuals give birth to new individuals at total per capita rate of *r*(*B*) and the locations of the new deprived individuals are sampled according to the density function of *B*. Eventually, satiated individuals become resource-deprived, and if they do not consume resources, they die:

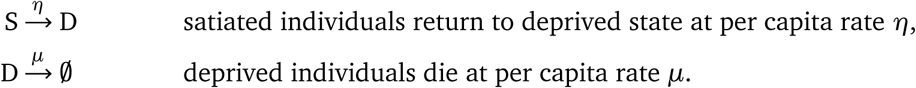

In addition to birth by resource-satiated individuals, we assume that there is a small background rate of influx immigration into the focal landscape:

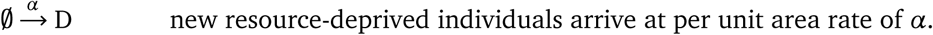

#### Deterministic, non-spatial model

Before we proceed, let us briefly check that the above resource-consumer model makes sense by considering the deterministic dynamics in a non-spatial setting, as this case is relatively simple to analyse (see Appendix A). In particular, a unique positive equilibrium of satiated individuals always exists if all rates are positive. In case *α* = 0, that is, when there is no immigration of individuals outside the focal landscape, the species goes extinct if the total resource production rate is not sufficiently high. More precisely, there exists an extinction threshold of

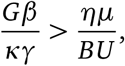

where *G*, *B*, *U* correspond to the total rates (i.e. integrals) of the respective kernels. Thus, the meanfield model already exhibits non-trivial single-species population dynamics. Now let us return to the stochastic, spatially-explicit and individual-based model with heterogeneous habitat structure.

#### Dispersal modes

We consider two modes of dispersal for the species:

- *passive* (one-shot) dispersal, where an individual does not move during its life time,
- *active* dispersal, where resource-deprived individuals move to find suitable habitat.

For passive dispersal, individuals only move when they are born according to the birth kernel *B* centered around their progenitor. In active dispersal, we include an additional movement process for resourced-deprived individuals:

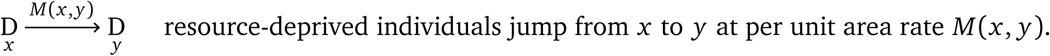

The idea is that in the active mode of dispersal, resource-deprived individuals can make additional movements to move from a location lacking resources to a new location with resources. Once an individual finds resources and consumes them, it becomes resource-satiated and stops moving.

We control the length of dispersal with the scale parameter *δ* and the mode of dispersal with an integer parameter *k* > 0. We set length scales of the Gaussian birth and movement kernels to 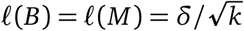 and set the movement rate to *r*(*M*) = (*k* – 1)*μ*. Passive dispersal corresponds to case *k =* 1, as no movement after birth occurs. In active dispersal, we have *k* > 1. Thus, species with passive dispersal are sessile, whereas species with active dispersal are not.

Note that regardless of the value *k*, the new location of an actively dispersing resource-deprived individual during its lifetime has the same mean and variance as a passive disperser *assuming the individual does not become satiated at any point in its lifetime.* If an individual remains resource-deprived its entire lifetime, it’s expected lifetime is 1/*μ* time units. This yields that the number of movement steps after birth is a Poisson random variable *s* with an expectation of *r* (*M*)/*μ* = *k* – 1. Hence, given the number of movement steps *s* (note that an individual always make a single movement step at birth), the total displacement is a random variable *x* = (*x*_1_, *x*_2_) that satisfies

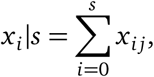

where *i* ∈ {1,2} and *x_ij_* ~ 𝒩(0,*δ*^2^/*k*) are independent Gaussian variables. Thus, *x_i_* follows a compound Poisson distribution with mean 0 and variance *δ*^2^ (i.e. same mean and variance as in the passive dispersal).

#### Habitat fragmentation

Above we have described the model in a pristine landscape with no habitat loss or fragmentation; while the habitat quality (resource production rate) can have patchy structure, the spatiotemporal dynamics of the resource patches guarantee that every location of the landscape has statistically the same properties over time.

We conduct our simulations in a finite landscape, and to avoid boundary effects, we assume it to be a two-dimensional torus ℒ ⊆ ℝ^2^ of size *L* × *L*. In order to incorporate habitat loss and fragmentation into our model, we assume that the focal landscape ℒ is partitioned into *N* disjoint circular *habitat fragments*, where the habitat fragment *i* is centered at location *x_i_*, has radius *r_i_*, and therefore, contains all points in the set

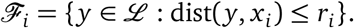

The *matrix* 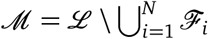 consists of the points that are not part of any habitat fragment.

We generate fragmented landscapes where the patch sizes follow a log-normal distribution so that there are patches of various sizes. Given the total habitat cover *C* and number of fragments *N*, we sample the relative area of each fragment *A_i_* ~ exp𝒩(*μ*, *σ*^2^) with parameters *μ* = 1/2 and *σ* = 1 for every fragment *i* ∈ {1,…,*N*} and normalise the areas so that ∑*A_i_* = *C* · *L*^2^.

The fragments are placed onto the landscape in a decreasing order in area and both coordinates of the fragment *i* are sampled uniformly at random from the interval [0, *L*], where *L* is the parameter controlling the size of the focal landscape. The coordinates of fragment *i* are resampled until there is no overlap with any fragment *j* < *i* to ensure that all fragments are disjoint. Figure 1 illustrates examples of fragmented landscapes for different values of *C* and *N*.

#### Effects of fragmentation

So far we have described how the fragmentation pattern is generated, but not how the pattern influences the system. We consider two different, extreme scenarios:

- *Habitable matrix*: Fragmentation only influences the resource production, but the species themselves are not (directly) affected by the matrix in any way. More precisely, resource particles can only establish within fragments, and to this end, Eq. **??** is modified so that a resource particle can only appear to location *y* if *y* ∉ ℳ. Any resource particle that falls into the matrix is immediately destroyed. The fragment and matrix have no other direct effects.
- *Hostile matrix*: Fragmentation influences directly both the resource production and the species survival. In addition to the above constraint on resource production, we assume that the species individuals cannot survive in the matrix. That is, any resource-deprived individual that immigrates or disperses into location *y* ∈ ℳ in the matrix is immediately killed. As satiated individuals do not move, it follows that satiated individuals can only reside within fragments.

The idea is that in the habitable matrix scenario makes minimal assumptions on effects of fragmentation, as it only affects how resource particles are generated. In particular, this scenario allows for *edge effects* to emerge (i.e. species can also survive in the matrix given close enough proximity to habitat fragments), whereas the hostile matrix scenario explicitly prevents this.

### 2.2 Simulation experiments

We simulated the model in a continuous torus landscape of size *L* × *L* with *L* = 100 (i.e. periodic boundary conditions) and *S* = 32 species all competing on the same limiting resource. Unless otherwise specified, the parameters are as in Table 1. In addition to the intact landscape (*N* = 1 and *C* = 1.0), we considered landscapes with varying number *N* of fragments and the fraction *C* of total area covered by the fragments; see Figure 2 for examples of the emerging community patterns.

**Table 1:**
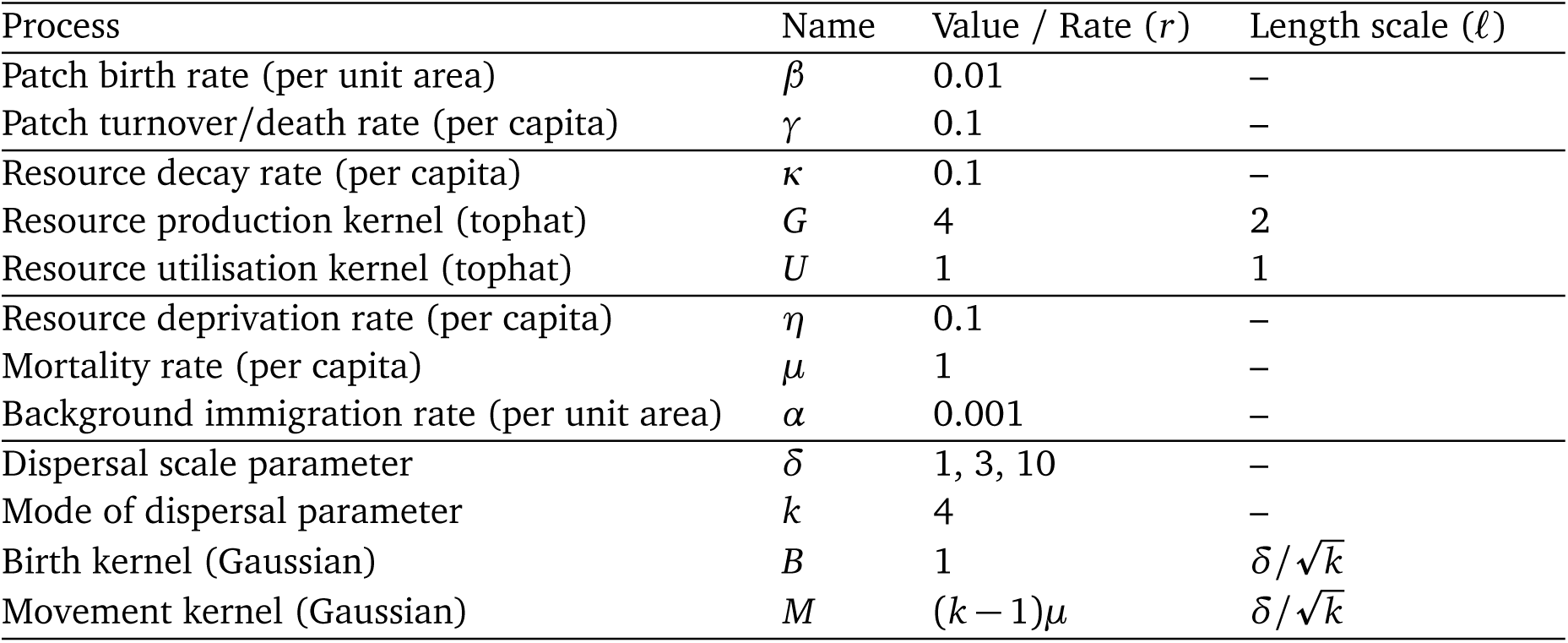
Parameters of the community model and their default values.

| Process | Name | Value / Rate ($r$) | Length scale ($\ell$) |
| --- | --- | --- | --- |
| Patch birth rate (per unit area) | $\beta$ | 0.01 | – |
| Patch turnover/death rate (per capita) | $\gamma$ | 0.1 | – |
| Resource decay rate (per capita) | $\kappa$ | 0.1 | – |
| Resource production kernel (tophat) | $G$ | 4 | 2 |
| Resource utilisation kernel (tophat) | $U$ | 1 | 1 |
| Resource deprivation rate (per capita) | $\eta$ | 0.1 | – |
| Mortality rate (per capita) | $\mu$ | 1 | – |
| Background immigration rate (per unit area) | $\alpha$ | 0.001 | – |
| Dispersal scale parameter | $\delta$ | 1, 3, 10 | – |
| Mode of dispersal parameter | $k$ | 4 | – |
| Birth kernel (Gaussian) | $B$ | 1 | $\delta/\sqrt{k}$ |
| Movement kernel (Gaussian) | $M$ | $(k-1)\mu$ | $\delta/\sqrt{k}$ |

**Figure 2:**
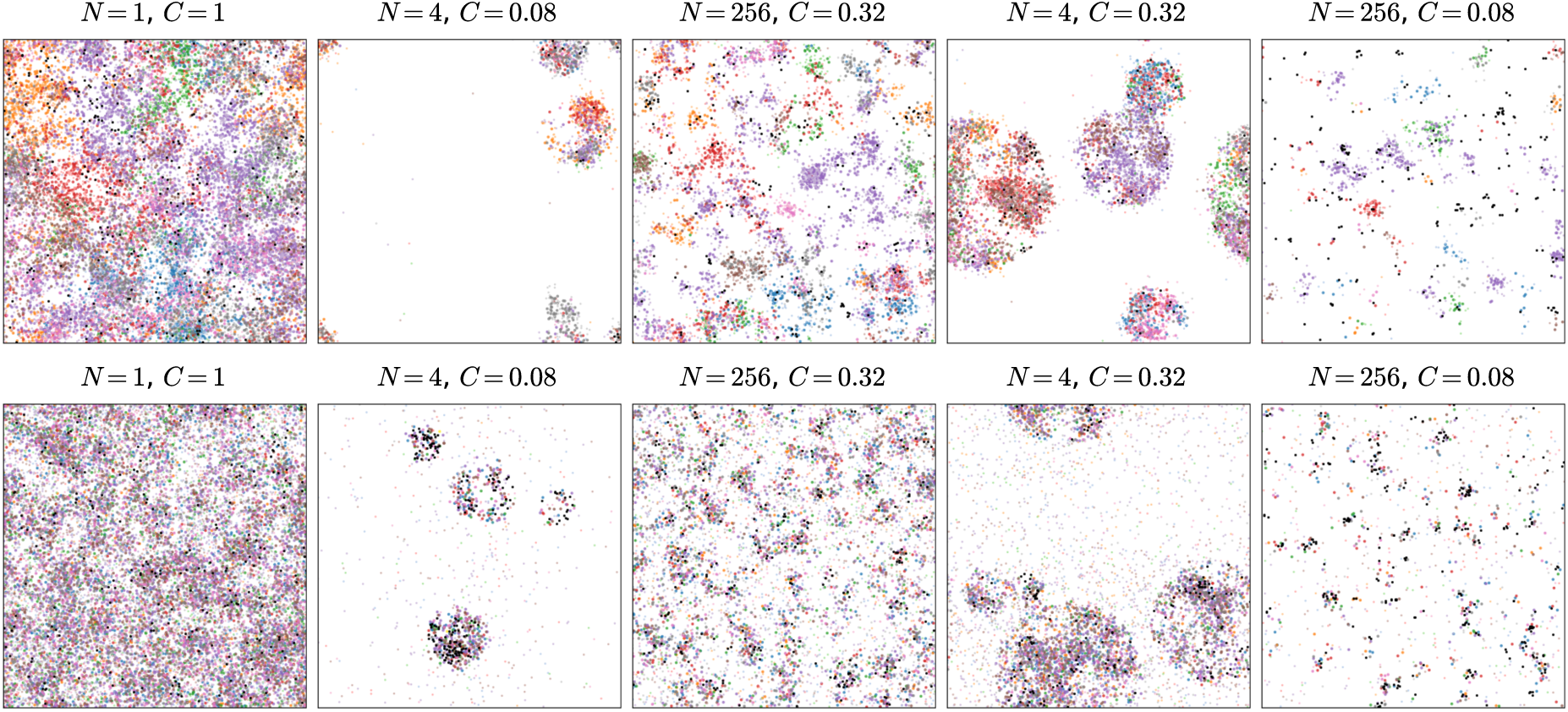
Examples snapshots of simulations of 8 species in landscapes with varying degree of fragmentation and habitat cover in a 50 × 50 landscape. The top row illustrates a scenario with *δ* = 1 and bottom row a scenario with *δ* = 10, both with passive dispersal. Coloured dots represent species individuals (larger ones are resource-satiated and smaller ones are resource-deprived individuals, respectively) and black dots represent resource particles. The resource patches and fragments are not drawn. In both cases the matrix is habitable. All other parameters are as in Table 1. The online supplementary material contains animated versions of the illustrated scenarios.

For each combination of number of patches *N* ∈{1,4,16,64,256,1024} and total habitat cover *C* ∈ {0.001,0.005,0.01,0.02,0.04,0.08,0.16,0.32}, we ran *R* = 100 replicate landscapes yielding a total of 6 × 8 × 100 = 4800 landscapes. For each replicate, we collected data of the total number of *resource*-*satiated* individuals (of each species) in

1. each fragment ℱ_*i*_,
2. the entire focal landscape ℒ, and
3. fixed-size sampling windows of radius *τ* = 1 centered in each fragment with radius of at least *τ*.

From data (1) we carried out the SLOSS (single large or several small) analysis (Fahrig, 2013), data (2) was used to plot SFAR (species fragmented-area relationship) curves (Rybicki and Hanski, 2013), and data from (3) was used to test the sample size effect as proposed by Fahrig (2013) and to examine community composition. The respective analyses were performed as follows.

#### SLOSS analysis

For the SLOSS curves, we sorted the fragments in both increasing and decreasing order and then plotted the cumulative number of species against the cumulative habitat amount. We then took the average of the cumulative species number *S_k_* and fraction of total habitat amount *A_k_* over all replicates, where

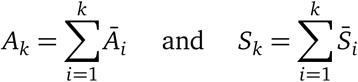

and *Ā_t_* and *S̄_i_* denote the average area and species count of the *i*th fragment in the given sorted order (either decreasing or increasing in area).

#### SFAR curves

For the SFAR curves, we plotted the average number of species *S* present in each simulated landscape. For every landscape with a total habitat cover of *C* and *N* fragments, we calculated the average number of species present in the entire landscape.

#### Testing the sample size effect

From the data (3), we calculated the average number of species as a function of the total habitat cover in the landscape and the area of the fragment in which the sample site was placed into.

#### Examining the community composition

To examine the community composition of species in a single replicate landscape, we computed *β*-diversity (community similarity) indices using data (3); we excluded landscapes with less than four sample sites. For each fixed-size sample site *i*, calculated the proportion **p**_*k*_ = (*p*_*k*,1_,…,*p*_*k*,*i*_,…, *p*_*k*,*S*_) of each species *i* in the sample. For the proportional data, we computed the average dissimilarity of species composition

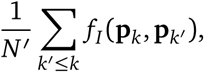

where *N*′ ≤ *N* is the number of fixed-size sample sites (one in each fragment that is large enough to fit the fixed-size sample site) and *f_I_*(**p**_*i*_, **p**_*i*′_) denotes the similarity of sites *i* and *i*′ according to index *I*. We examined two similarity indices: the generalised Jaccard index

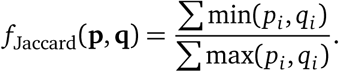

and the Euclidean distance

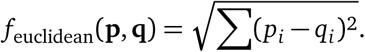

## 3 Results

We set out to test how fragmentation influences species richness under different conditions. We considered three scales of dispersal (from short *δ* = 1, to intermediate *δ* = 3 and long-range *δ* = 10), two modes of dispersal (passive and active), and two matrix types (habitable and hostile). For the sake of simplicity, we excluded the scenario with active dispersal and hostile matrix, as any individual moving outside the habitat fragment would immediately die in a hostile matrix, thus limiting the potential benefit of active dispersal under fragmentation. In total we obtained 9 different types of species communities, each of which was simulated in total of 4800 replicate landsacpes (i.e. with different fragmentation patterns).

### 3.1 The SLOSS analysis

In order to disentangle the effect of habitat loss and fragmentation, Fahrig (2013) suggested conducting SLOSS analyses. As discussed, for the SLOSS curves, the fragments are sorted in both increasing and decreasing order of area, and then the cumulative number of species is plotted against the cumulative habitat amount. As this needs to be done for each landscape separately, this yields thousands of plots for each of the nine community scenarios which we have considered.

The result of all these analyses is summarised in Figure 3. We can see that when the dispersal distances are short or intermediate, fragmentation shows both positive and negative effects. This depends on the total amount of habitat available: at low habitat cover, fragmentation is detrimental, whereas fragmentation in landscapes with high habitat cover has a positive effect.

**Figure 3:**
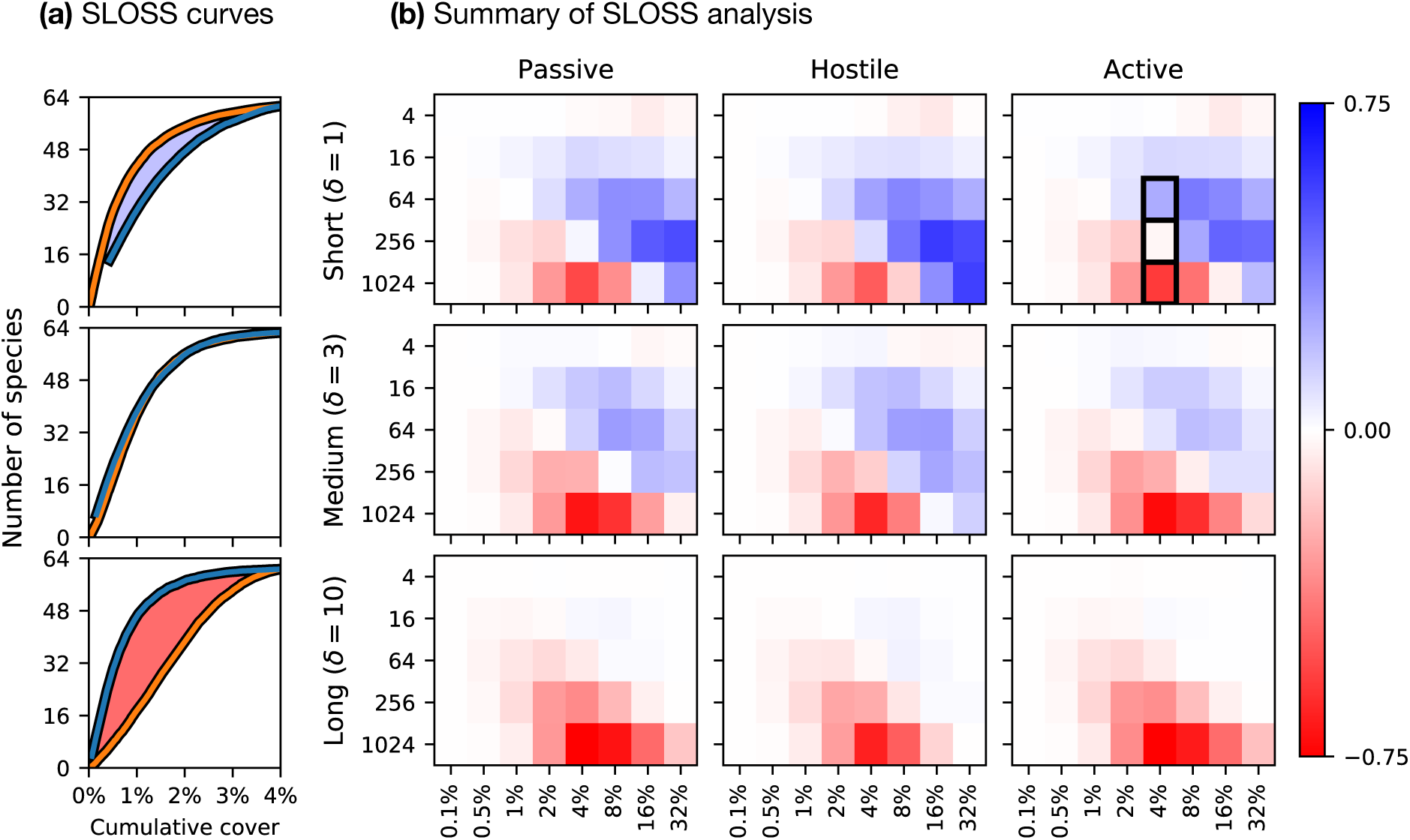
Summary of the SLOSS analysis. (a) Example SLOSS analysis curves. Here, the habitat fragments have been sorted in increasing (from smallest patch to largest, orange) and decreasing (largest patch to the smallest, blue) order. The horizontal axis indicates the *cumulative* habitat cover and the vertical axis the *cumulative* number of species averaged over all replicates. When the orange line is above the blue line, fragmentation has a positive effect on species richness (top box), whereas when the blue line is above the orange line, fragmentation has a negative effect on species richness (bottom box). If both lines overlap (middle box), then fragmentation *per se* has no effect (as predicted by the habitat amount hypothesis). (b) For each value of *δ* and dispersal mode, there is a coloured grid that summarises the SLOSS analysis for 5 × 8 landscape scenarios. In each grid, the vertical axis gives the total number of fragments in the landscape and the horizontal axis the total habitat cover. The colour of the cell indicates the *area* of the integral between the orange and blue lines. Blue indicates positive effect of fragmentation (orange line above blue line, top box) and red indicates negative effect (blue above orange line, bottom box). Thus, stronger the colour, more pronounced the effect of fragmentation. The example plots in (a) are given by the highlighted rectangles in the third grid of the top row (*δ* = 1 with active dispersal).

### 3.2 Testing the sample size effect

For testing the sample size effect, Fahrig (2013) suggested examining the number of species in fixed-size sample sites, as a function of the area of in the local landscape. Figure 4 shows the average number of species in fixed-size sampling sites as a function of the total landscape cover.

**Figure 4:**
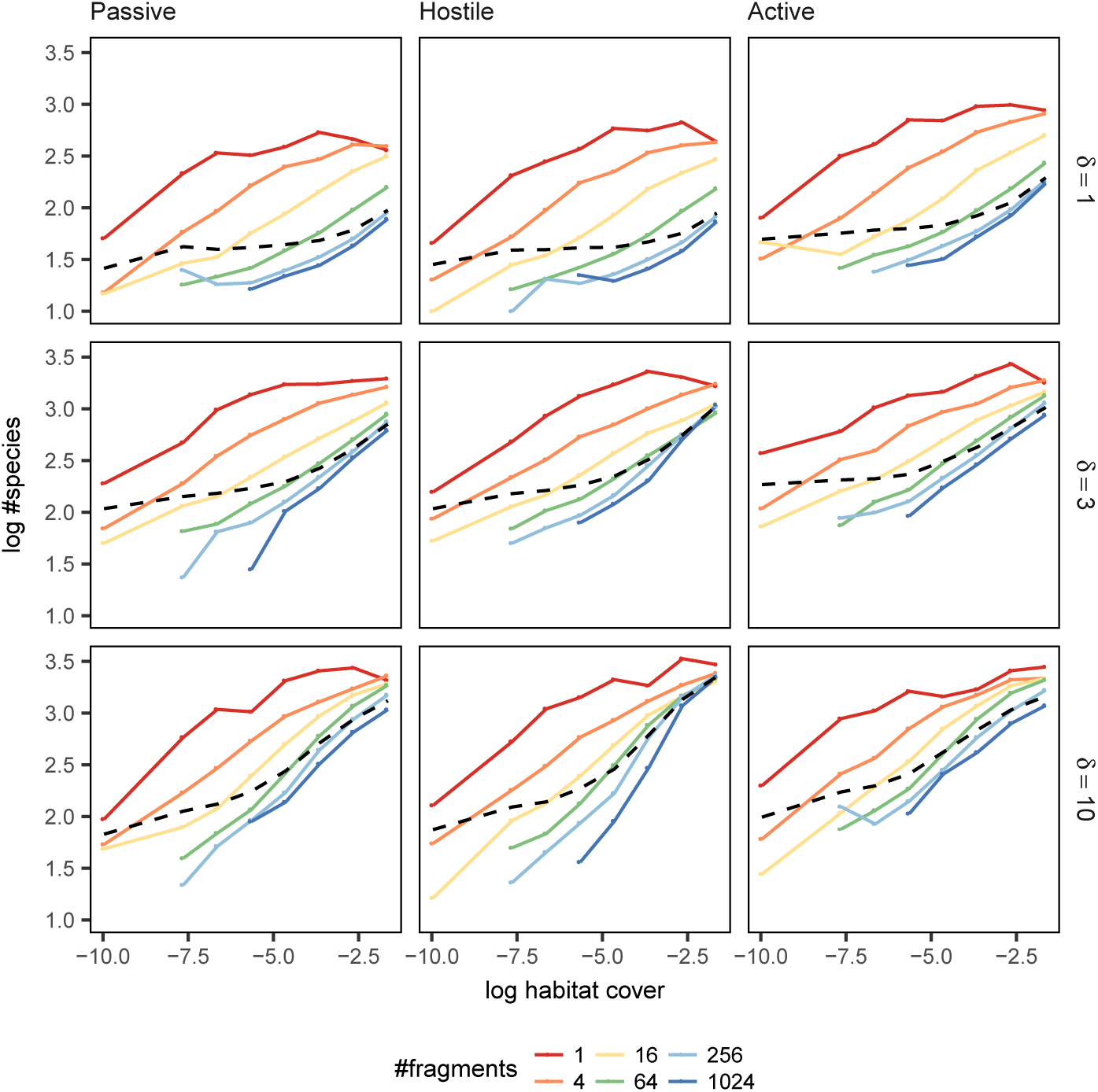
The average number of species in fixed-size sample sites of radius *τ* = 1 centered in each fragment with at least radius *τ*. In each plot, the horizontal axis indicates the average habitat cover of the landscape and vertical axis the average number of species in a fixed-size sample site (data points with less than five replicates were omitted). Each coloured line shows data from landscapes with a different numbers of fragments with the exception of the black dotted line which shows the average over all landscapes. The different rows show community scenarios with increasing dispersal range from top to bottom, whereas the columns represent different dispersal/matrix scenarios.

Given the same amount of total habitat, in all cases the sample sites contained more species when the surrounding landscape was divided into fewer fragments. In order to be consistent with the habitat amount hypothesis, the different lines should be indistinguishable, as only the total amount of habitat should matter. However, it can be seen that the total of number fragments in the landscape, and thus the landscape structure, clearly influences the number of species at fixed-size sample sites.

When looking at the average number of species in fixed-size sample sites over all landscapes (black dashed line in Figure 4), the line representing the relationship is almost horizontal when the total habitat amount is low or the dispersal scale is short. This indicates that the average species richness is little influenced by the total habitat cover. However, when the total amount of habitat is large, there is a strong positive relationship with the number of species and habitat cover. Thus, when the habitat cover is low and dispersal distances short, the results are inconsistent with the habitat amount hypothesis. However, if the dispersal distance is large and the habitat cover is high, the results become more consistent with the habitat amount hypothesis.

### 3.3 Species-area relationships

Whereas the SLOSS curves only account for species observed *within* habitat fragments, the SFAR curves (Figure 5) show the species diversity in the entire focal landscape (i.e. how many species were present in the landscape). Note that even if there is a single resource-satiated individual in the landscape, then the species is counted to be present (no matter how unlikely sampling it might be).

**Figure 5:**
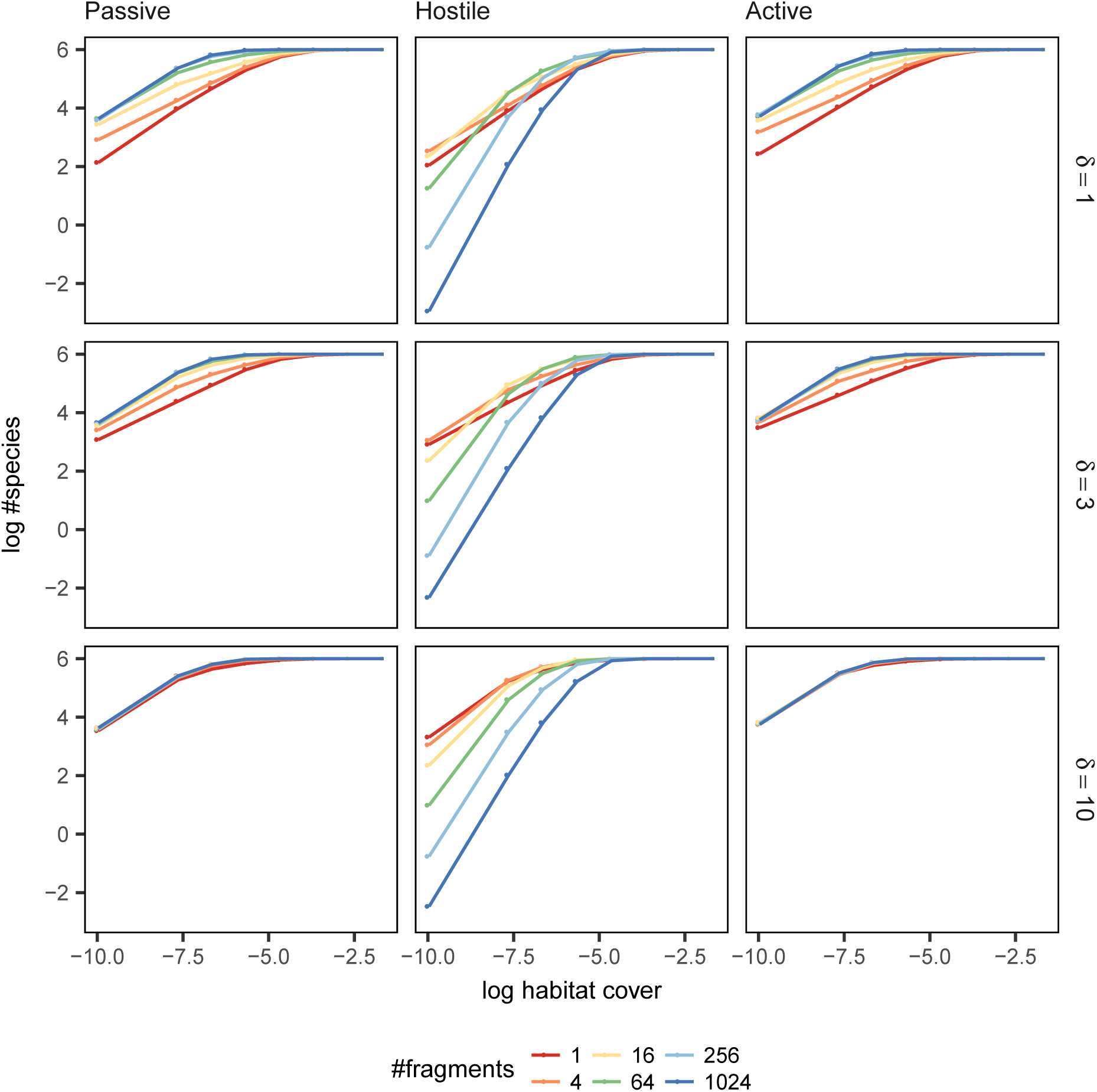
Species-fragmented area curves for different scenarios. The plot shows the average number of species (vertical axis) over in a landscape of given total habitat cover (horizontal axis) and number of fragments (averaged over all replicates). The rows represent results for different dispersal ranges and columns for different dispersal/matrix scenarios.

The species-area relationship exhibits different patterns depending on whether the matrix is habitable or hostile for the focal species. If the matrix is hostile, that is, the individuals cannot move and survive the matrix, then the SAR plots show higher levels of fragmentation being detrimental for species diversity. In contrast, if the matrix is habitable, that is, individuals can move and survive in the matrix (but the resources still have to be obtained from habitat fragments), the SAR suggests that species richness only depends only on the total amount of habitat in the landscape.

### 3.4 Community composition

Recall that *β*-diversity measures the species turnover, that is, variability of species composition across different communities. Figure 6 shows how the average similarity in community composition behaves under the different scenarios. We see that habitat loss increases dissimilarity and increasing dispersal distance decreases dissimilarity. Moreover, we see that the average dissimilarity slightly increases with increasing fragmentation (i.e. number of fragments in the landscape). Thus, given the same amount of total habitat, *β*-diversity increases with increasing dispersal distance and level of fragmentation. This is expected, as stochastic events and competitive exclusion will drive some species out locally. The same observations hold also when using larger sample sites with radius *τ* = 2 (but we see greater variance in similarity, see Figure 8), and when using the Euclidean distance as a similarity measure (Figure 9, Figure 10).

**Figure 6:**
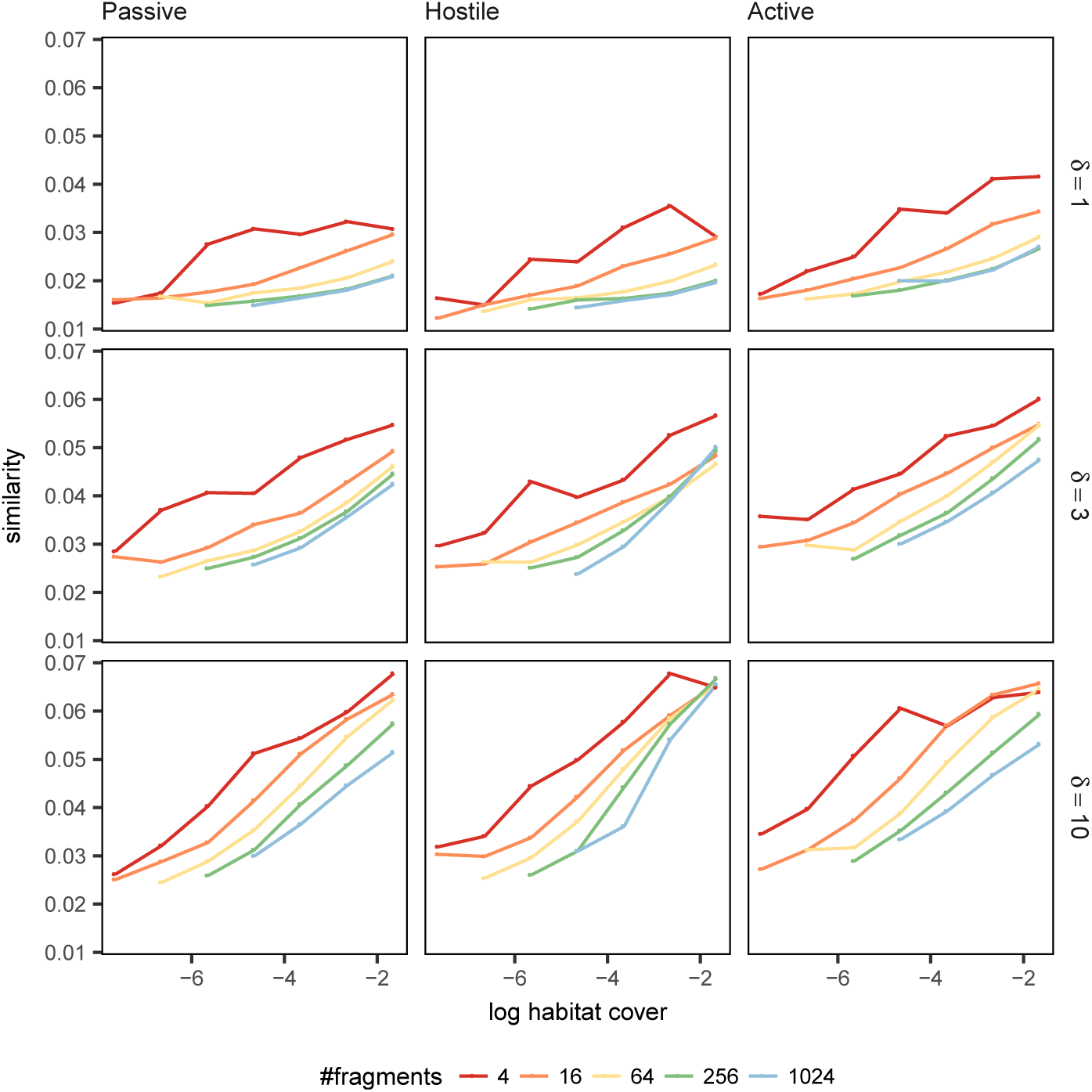
The average Jaccard similarity in community composition on fixed-size sample sites of radius *τ* = 1. The *x*-axis gives the logarithm of the total habitat cover in a landscape, and the *y*-axis gives the average similarity of sample sites (averaged over all landscape replicates). Higher similarity values on *y*-axis indicate that the species composition is on average more similar in landscapes with given habitat cover.

### 3.5 Summary of results

We have examined how species richness and diversity is influenced by habitat loss and fragmentation in different competitive species communities; essentially, we have measured

- local species richness at sample sites (*α*-diversity),
- species turnover across different sample sites (*β*-diversity), and
- total species diversity in a landscape (*γ*-diversity).

In summary, we observe that with *increasing fragmentation* the situation differs whether the total amount of habitat is high or low. With *high* amount of total habitat, fragmentation

- can show positive effects in the SLOSS analysis, especially when dispersal distances are short,
- slightly reduces species richness at fixed-size sample sites (*α*-diversity),
- can slightly increase species turnover (*β*-diversity),
- has little effect at landscape level, that is, species richness is mostly unaffected (*γ*-diversity).

That is, fragmentation *per se* has either no significant negative effect or some positive effects given that the total amount of habitat remains high in the landscape. Conversely, if the total amount of habitat is *low*, then fragmentation

- consistently yields negative effects according to the SLOSS analysis,
- reduces species richness at fixed-size sample sites (*α*-diversity),
- can slightly increase species turnover (*β*-diversity),
- greatly reduces species richness at the landscape level if the matrix is hostile, but can slightly increase species richness if the species can survive in the matrix (*γ*-diversity),

That is, fragmentation *per se* (dividing the same amount of total habitat into smaller pieces) has a negative effect on species richness if the total amount of habitat is low, but in terms of community composition, increased fragmentation can slightly increase *β*-diversity.

## 4 Discussion

The question of how landscape structure and fragmentation *per se* affect species richness has been long debated. The habitat amount hypothesis posits that fragmentation *per se* does not have a strong effect, but it is only the total amount of habitat in a local landscape that matters (Fahrig, 2013), and a metanalysis of empirical fragmentation studies shows that fragmentation can have negative and positive effects on species richness (Fahrig, 2017). However, the habitat amount hypothesis has been criticised for lacking a mechanistic explanation (Hanski, 2015). In this work, we examined when does the habitat amount hypothesis hold following the tests outlined by Fahrig (2013) using a mechanistic individual-based metacommunity model of habitat specialists.

Our results suggest that, given the same total amount of habitat, fragmentation *per se* can show *both* negative and positive effects on species richness in competitive metacommunities. If the total amount of habitat is small, then fragmentation *per se* negatively affects species richness. On the other hand, if the total amount of habitat is sufficiently large, then fragmentation can show a positive effect on species richness of competing habitat specialists (e.g. Figure 3). However, in various intermediate scenarios fragmentation exhibits little effect, which suggests that the habitat amount hypothesis may be a reasonable approximation in some circumstances. Overall, different analyses on the same data can seemingly exhibit different responses to fragmentation (Figures 3–5). This highlights that caution is needed when analysing the effects of fragmentation and may be one reason as to why fragmentation has been reported to have so varying effects on species richness (With, 2016; Fahrig, 2017; Arnillas et al., 2017; Haddad et al., 2017; De Camargo et al., 2018).

In general, our results are inconsistent with the habitat amount hypothesis – at least when the total amount of habitat is small – which suggests that either (1) the habitat amount hypothesis does not hold in all situations, (2) the hypothesis needs to be refined, or (3) our model lacks some critical features. Of course, all models are simplifications of reality and omit various details. Indeed, prior models have been critisised, e.g., De Camargo et al. (2018) suggest that “model of area-dependent, stochastic patch occupancy and extinction” by Rybicki and Hanski (2013) “fails to capture some critical aspect of habitat loss”, and hence, exhibit adverse effects of fragmentation. Nevertheless, in this work we have used a model with fundamentally different structural assumptions and yet still obtain the same qualitative results, and our results are consistent with prior theory and modelling work (Tilman et al., 1997; Rybicki and Hanski, 2013).

In any case, whether or not our model captures all key aspects governing species dynamics under fragmentation, mathematical models can still help shed light on which mechanisms drive the community patterns emerging in fragmented landscapes. Thus, even if the habitat amount hypothesis is true, the underlying mechanisms are poorly understood. Our work suggests that the hypothesis should rely on some mechanisms not accounted in our or any of the prior models, e.g. fast eco-evolutionary responses (Legrand et al., 2017). Furthermore, not all species are equally sensitive to habitat fragmentation and how different species tolerate the matrix varies (Gascon et al., 1999). In this work, we have restricted our attention to competitive habitat specialists, and thus, areas inflicted with habitat loss cannot sustain any species (unless near habitat). In particular, we do not consider *habitat conversion*, where one type of habitat is converted to another, which may still be suitable to some existing species or new species that can replace the original species in the species pool.

Beyond examining the effects of fragmentation effects on species richness, we believe that our modelling framework lends itself to be a useful tool for spatially-explicit investigations of various other species interactions in metacommunities. So far, our work only considers communities of competing habitat specialists, and thus, it could be extended to other community structures and dynamics, such as predator-prey dynamics, competition-colonization tradeoffs, mutualist species, or successive metacommunities. Such extensions may reveal how responses to fragmentation depend on the specific of metacommunity dynamics. Indeed, Wilson et al. (2016) point out that “other measures of community structure, such as community composition, trophic organization, species persistence, and species residency, may better inform how fragmentation affects biotic communities, even when species richness per se is not altered by fragmentation”.

## Acknowledgements

The authors wish to acknowledge CSC — IT Center for Science, Finland, for computational resources. This project has received funding from the European Union’s Horizon 2020 research and innovation programme under the Marie Skłodowska-Curie grant agreement No. 754411 (J.R.) and from the Academy of Finland postdoctoral grant 308651 (N.A.).

## A Meanfield approximation of species dynamics

Let us examine the single-species dynamics in a non-spatial setting. If we assume that the system is well-mixed and let the length scale *ℓ* → ∞ for each kernel, then the deterministic meanfield model can be written as

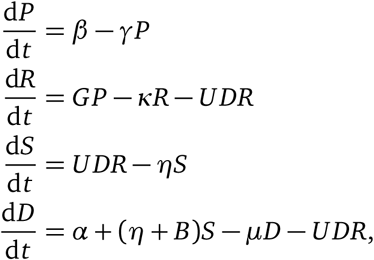

where the capital letter *X* denotes the integral *r*(*X*) of the kernel *X*, *a* is non-negative and all other parameters are positive. The two non-trivial equilibria are

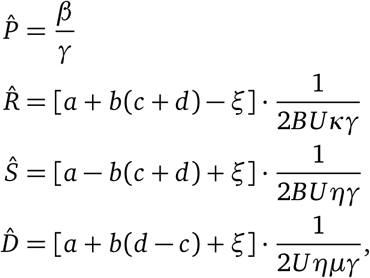

where we abbreviate

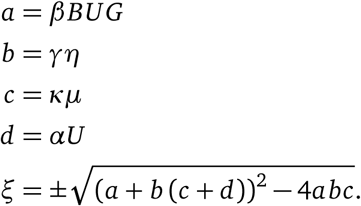

Since we have *a*, *b*, *c* > 0 and *d* ≥ 0, this yields that *ξ* is real, as

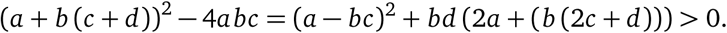

Therefore, a unique positive equilibrium *Ŝ* > 0 of satiated individuals exists when

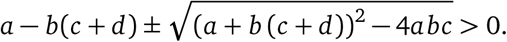

If the third term has a negative sign, the condition cannot be satisfied. However, if the third term has a positive sign, then the condition is always satisfied if all parameters *a*, *b*, *c*, *d* > 0 are positive. In the case that *α* = 0, that is, there is no immigration of individuals from outside the focal landscape, we have *d* = 0 and a positive equilibrium exists only if *a* > *bc* holds. Rewriting this condition gives

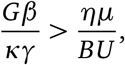

where the left-hand side corresponds to the resource density of the habitat in the absence of the species. Thus, without outside immigration, the species cannot persist if the total amount and/or quality of habitat is not sufficiently high. In other words, we obtain an extinction threshold for the deterministic, non-spatial single-species model in the absence of outside immigration.

## B Supplementary material

### B.1 Online material

Further supplementary material is available online^∗^. The online supplementary material contains

- Source code to the simulator and the analysis scripts, and
- additional animations of the scenarios illustrated in Figure 2.

### B.2 Additional illustrations

Below we have included the following supplementary illustrations:

- Figure 7 shows the analysis of Figure 4 for the case *τ* = 2,
- Figure 8 shows the analysis of Figure 6 for the case *τ* = 2, and
- Figure 9 and Figure 10 show the community composition analysis (Figure 6) using Euclidean distance as a similarity metric for *τ* = 1 and *τ* = 2, respectively.

**Figure 7:**
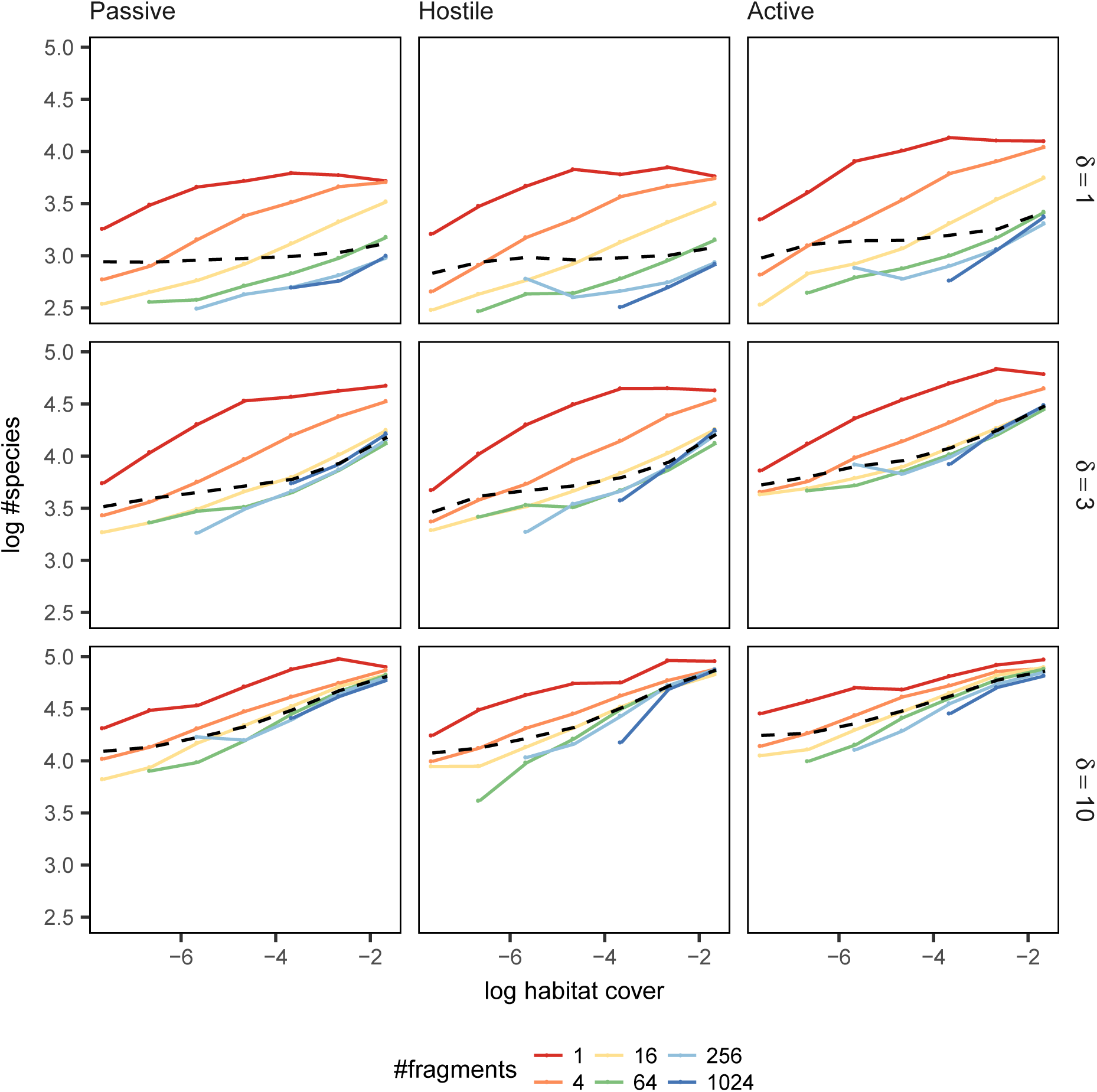
The average number of species in fixed-size sample sites of radius *τ* = 2 centered in each fragment with at least radius *τ* (see Figure 4 for the case *T* = 1). In each plot, the horizontal axis indicates the average habitat cover of the landscape and vertical axis the average number of species in a fixed-size sample site (data points with less than five replicates were omitted). Each coloured line shows data from landscapes with a different numbers of fragments with the exception of the black dotted line which shows the average over all landscapes. The different rows show community scenarios with increasing dispersal range from top to bottom, whereas the columns represent different dispersal/matrix scenarios.

**Figure 8:**
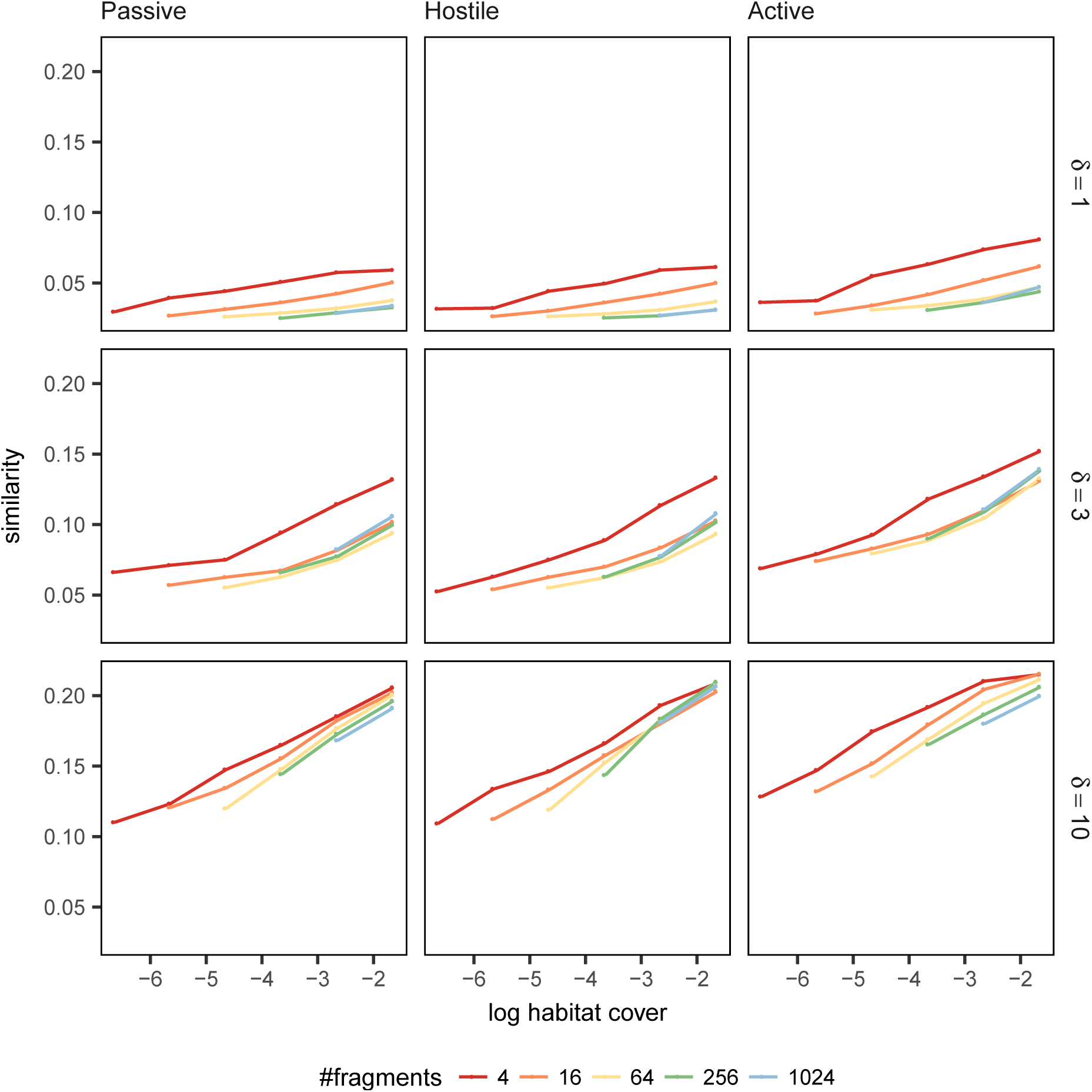
The average Jaccard similarity in community composition on fixed-size sample sites of radius *τ* = 2 (see Figure 6 for the case *τ* = 1). The *x*-axis gives the logarithm of the total habitat cover in a landscape, and the *y*-axis gives the average similarity of sample sites (averaged over all landscape replicates). Higher similarity values on *y*-axis indicate that the species composition is on average more similar in landscapes with given habitat cover.

**Figure 9:**
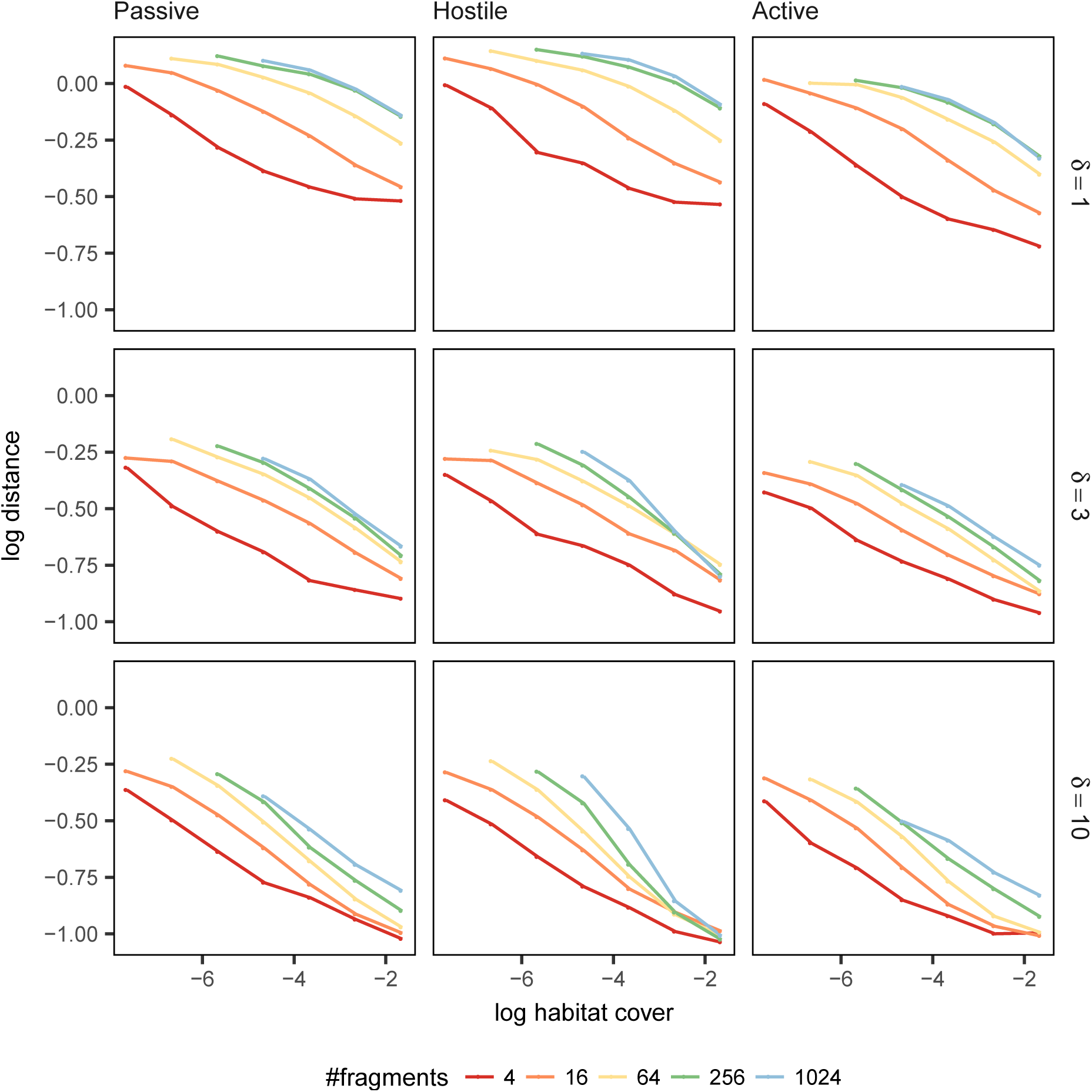
The average log Euclidean distance in community composition on fixed-size sample sites of radius *τ* = 1. The *x*-axis gives the logarithm of the total habitat cover in a landscape, and the *y*-axis gives the logarithm of the average Euclidean distance of the vectors describing the community composition at sample sites (averaged over all replicates). Lower values on the *y*-axis indicate that the species composition is on average more similar in landscapes with given habitat cover.

**Figure 10:**
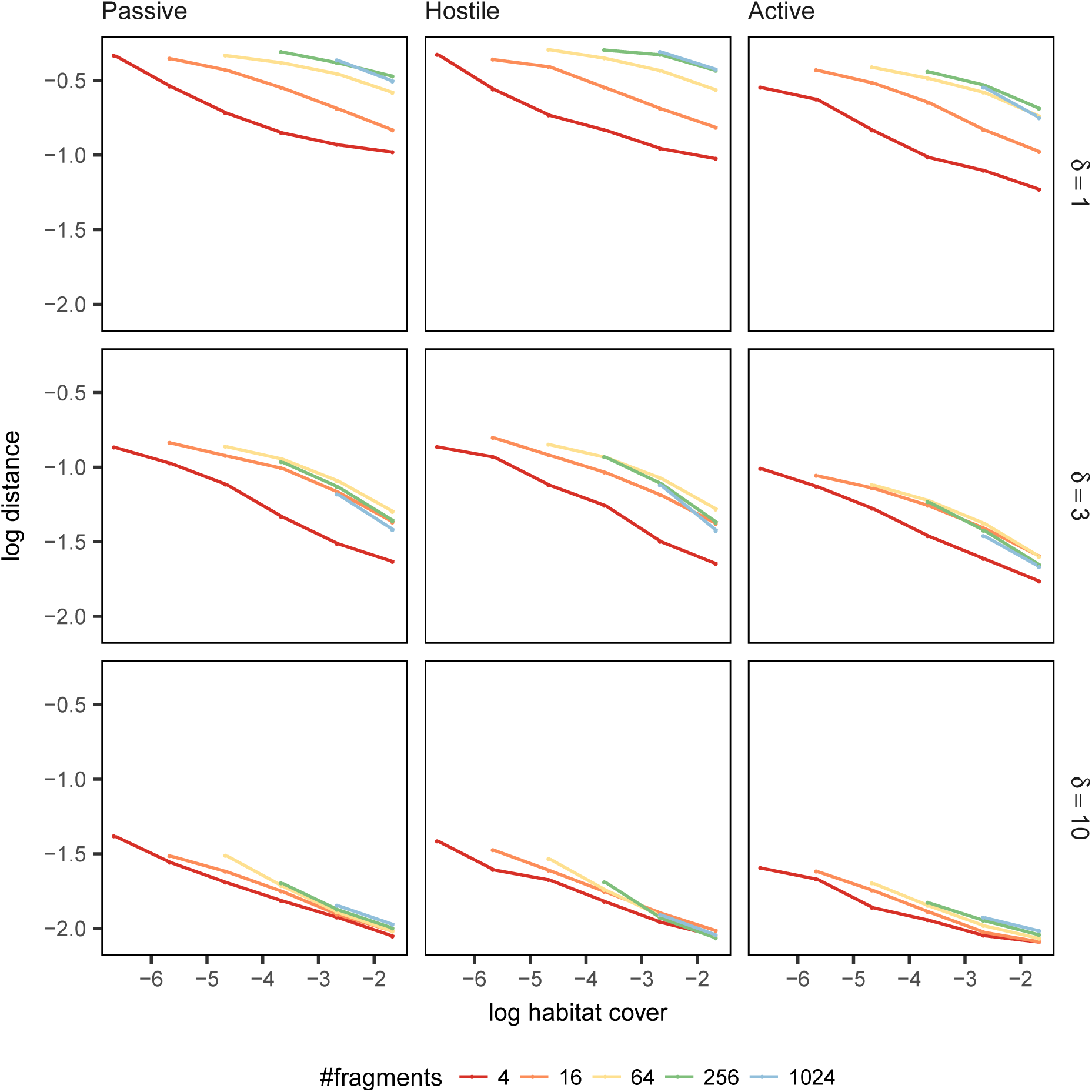
The average log Euclidean distance in community composition on fixed-size sample sites of radius *τ* = 2. The *x*-axis gives the logarithm of the total habitat cover in a landscape, and the *y*-axis gives the logarithm of the average Euclidean distance of the vectors describing the community composition at sample sites (averaged over all replicates). Lower values on the *y*-axis indicate that the species composition is on average more similar in landscapes with given habitat cover.

## Footnotes

∗ Online supplementary material: https://bitbucket.org/jrybicki/habitat-fragmentation

∗ https://bitbucket.org/jrybicki/habitat-fragmentation

## References

Carlos Alberto Arnillas, Carolina Tovar, Marc W. Cadotte, and Wouter Buytaert. From patches to richness: assessing the potential impact of landscape transformation on biodiversity. Ecosphere, 8 (11):e02004, 2017. doi:10.1002/ecs2.2004.

Jordi Bascompte and Ricard V Solé. Habitat fragmentation and extinction thresholds in spatially explicit models. The Journal of Animal Ecology, 65(4):465, 1996. doi:10.2307/5781.

Stuart H. M. Butchart, Matt Walpole, Ben Collen, Arco van Strien, Jörn P W Scharlemann, Rosamunde E. A. Almond, Jonathan E. M. Baillie, Bastian Bomhard, Claire Brown, John Bruno, Kent E. Carpenter, Geneviève M. Carr, Janice Chanson, Anna M. Chenery, Jorge Csirke, Nick C. Davidson, Frank Dentener, Matt Foster, Alessandro Galli, James N. Galloway, Piero Genovesi, Richard D. Gregory, Marc Hockings, Valerie Kapos, J.-F. Lamarque, Fiona Leverington, Jonathan Loh, Melodie A McGeoch, Louise McRae, Anahit Minasyan, M. H. Morcillo, Thomasina E. E. Oldfield, Daniel Pauly, Suhel Quader, Carmen Revenga, John R. Sauer, Benjamin Skolnik, Dian Spear, Damon Stanwell-Smith, Simon N. Stuart, Andy Symes, Megan Tierney, Tristan D. Tyrrell, J.-C. Vie, and Reg Watson. Global biodiversity: Indicators of recent declines. Science, 328(5982):1164–1168, 2010. doi:10.1126/science.1187512.

Rafael X. De Camargo, Véronique Boucher-Lalonde, and David J. Currie. At the landscape level, birds respond strongly to habitat amount but weakly to fragmentation. Diversity and Distributions, 2018. doi:10.1111/ddi.12706.

Jared M. Diamond. The island dilemma: Lessons of modern biogeographic studies for the design of natural reserves. Biological Conservation, 7(2):129–146, 1975. doi:10.1016/0006-3207(75)90052-X.

Raphael K. Didham, Valerie Kapos, and Robert M. Ewers. Rethinking the conceptual foundations of habitat fragmentation research. Oikos, 121(2):161–170, 2012. doi:10.1111/j.1600-0706.2011.20273.x.

Robert M. Ewers and Raphael K. Didham. Confounding factors in the detection of species responses to habitat fragmentation. Biological Reviews, 81(01):117, 2005. doi:10.1017/S1464793105006949.

Lenore Fahrig. Effects of habitat fragmentation on biodiversity. Annual Review of Ecology, Evolution, and Systematics, 34(1):487–515, 2003. doi:10.1146/annurev.ecolsys.34.011802.132419.

Lenore Fahrig. Rethinking patch size and isolation effects: The habitat amount hypothesis. Journal of Biogeography, 40(9):1649–1663, 2013. doi:10.1111/jbi.12130.

Lenore Fahrig. Just a hypothesis: a reply to Hanski. Journal of Biogeography, 42(5):993–994, 2015. doi:10.1111/jbi.12504.

Lenore Fahrig. Ecological responses to habitat fragmentation per se. Annual Review of Ecology, Evolution, and Systematics, 48(1):annurev–ecolsys–110316–022612, 2017. doi:10.1146/annurev-ecolsys-110316-022612.

Miguel A. Fortuna and Jordi Bascompte. Habitat loss and the structure of plant-animal mutualistic networks. Ecology Letters, 9(3):281–286, 2006. doi:10.1111/j.1461-0248.2005.00868.x.

Bertrand Fournier, Nicolas Mouquet, Mathew A. Leibold, and Dominique Gravel. An integrative framework of coexistence mechanisms in competitive metacommunities. Ecography, (April):1–12, 2016. doi:10.1111/ecog.02137.

Claude Gascon, Thomas E. Lovejoy, Richard O. Bierregaard Jr., Jay R. Malcolm, Phillip C. Stouffer, Heraldo L. Vasconcelos, William F. Laurance, Barbara Zimmerman, Mandy Tocher, and SÃľrgio Borges. Matrix habitat and species richness in tropical forest remnants. Biological Conservation, 91 (2):223–229, 1999. ISSN 0006-3207. doi:10.1016/S0006-3207(99)00080-4.

Luis J. Gilarranz and Jordi Bascompte. Spatial network structure and metapopulation persistence. Journal of Theoretical Biology, 297:11–16, 2012. doi:10.1016/J.JTBI.2011.11.027.

Dominique Gravel, Charles D. Canham, Marilou Beaudet, and Christian Messier. Reconciling niche and neutrality: the continuum hypothesis. Ecology Letters, 9(4):399–409, 2006. doi:10.1111/j. 1461-0248.2006.00884.x.

Dominique Gravel, François Massol, and Mathew A. Leibold. Stability and complexity in model meta-ecosystems. Nature Communications, 7(10):12457, 2016. doi:10.1038/ncomms12457.

Jacopo Grilli, György Barabás, and Stefano Allesina. Metapopulation persistence in random fragmented landscapes. PLOS Computational Biology, 11(5):e1004251, 2015. doi:10.1371/journal.pcbi.1004251.

N. M. Haddad, L. A. Brudvig, J. Clobert, K. F. Davies, A. Gonzalez, R. D. Holt, T. E. Lovejoy, J. O. Sexton, M. P. Austin, C. D. Collins, W. M. Cook, E. I. Damschen, R. M. Ewers, B. L. Foster, C. N. Jenkins, A. J. King, W. F. Laurance, D. J. Levey, C. R. Margules, B. A. Melbourne, A. O. Nicholls, J. L. Orrock, D.-X. Song, and J. R. Townshend. Habitat fragmentation and its lasting impact on earth’s ecosystems. Science Advances, 1(2):e1500052–e1500052, 2015. doi:10.1126/sciadv.1500052.

Nick M. Haddad, Andrew Gonzalez, Lars A. Brudvig, Melissa A. Burt, Douglas J. Levey, and Ellen I. Damschen. Experimental evidence does not support the habitat amount hypothesis. Ecography, 40 (1):48–55, 2017. doi:10.1111/ecog.02535.

Bart Haegeman and Michel Loreau. General relationships between consumer dispersal, resource dispersal and metacommunity diversity. Ecology Letters, 17(2):175–184, 2014. doi:10.1111/ele.12214.

Ilkka Hanski. Metapopulation Ecology. Oxford University Press, 1999.

Ilkka Hanski. Habitat fragmentation and species richness. Journal of Biogeography, 42(5):989–993, 2015. doi:10.1111/jbi.12478.

Ilkka Hanski and Otso Ovaskainen. The metapopulation capacity of a fragmented landscape. Nature, 404(6779):755–758, 2000. doi:10.1038/35008063.

Ilkka Hanski and Otso Ovaskainen. Metapopulation theory for fragmented landscapes. Theoretical Population Biology, 64(1):119–127, 2003. doi:10.1016/S0040-5809(03)00022-4.

Ilkka Hanski, Gustavo a Zurita, M Isabel Bellocq, and Joel Rybicki. Species-fragmented area relationship. Proceedings of the National Academy of Sciences, 110(31):12715–12720, 2013. doi:10.1073/pnas.1311491110.

Klaus Henle, Kendi F. Davies, Michael Kleyer, Chris Margules, and Josef Settele. Predictors of species sensitivity to fragmentation. Biodiversity & Conservation, 13(1):207–251, jan 2004. doi:10.1023/B: BIOC.0000004319.91643.9e.

Heather Bird Jackson and Lenore Fahrig. What size is a biologically relevant landscape? Landscape Ecology, 27(7):929–941, 2012. doi:10.1007/s10980-012-9757-9.

Aapo Kahilainen, Saskya Nouhuys, Torsti Schulz, and Marjo Saastamoinen. Metapopulation dynamics in a changing climate: Increasing spatial synchrony in weather conditions drives metapopulation synchrony of a butterfly inhabiting a fragmented landscape. Global Change Biology, 2018. doi:10.1111/gcb.14280.

C.A. Klausmeier. Habitat destruction and extinction in competitive and mutualistic metacommunities. Ecology Letters, 4(1):57–63, 2001. doi:10.1046/j.1461-0248.2001.00195.x.

Mikko Kuussaari, Riccardo Bommarco, Risto K Heikkinen, Aveliina Helm, Jochen Krauss, Regina Lindborg, Erik Öckinger, Meelis Pärtel, Joan Pino, Ferran Rodá, Constantí Stefanescu, Tiit Teder, Martin Zobel, and Ingolf Steffan-Dewenter. Extinction debt: a challenge for biodiversity conservation. Trends in Ecology & Evolution, 24(10):564–571, 2009. doi:10.1016/j.tree.2009.04.011.

Delphine Legrand, Julien Cote, Emanuel A. Fronhofer, Robert D. Holt, Ophélie Ronce, Nicolas Schtickzelle, Justin M. J. Travis, and Jean Clobert. Eco-evolutionary dynamics in fragmented landscapes. Ecography, 40(1):9–25, 2017. doi:10.1111/ecog.02537.

Clarence L. Lehman and David Tilman. Biodiversity, stability, and productivity in competitive communities. The American Naturalist, 156(5):534–552, 2000. doi:10.1086/303402.

M. A. Leibold, M. Holyoak, N. Mouquet, P. Amarasekare, J. M. Chase, M. F. Hoopes, R. D. Holt, J. B. Shurin, R. Law, D. Tilman, M. Loreau, and A. Gonzalez. The metacommunity concept: A framework for multi-scale community ecology. Ecology Letters, 7(7):601–613, 2004. doi:10.1111/j. 1461-0248.2004.00608.x.

Jürg B. Logue, Nicolas Mouquet, Hannes Peter, and Helmut Hillebrand. Empirical approaches to metacommunities: a review and comparison with theory. Trends in Ecology & Evolution, 26(9): 482–491, 2011. doi:10.1016/J.TREE.2011.04.009.

Michel Loreau, Nicolas Mouquet, and Andrew Gonzalez. Biodiversity as spatial insurance in heterogeneous landscapes. Proceedings of the National Academy of Sciences of the United States of America, 100(22):12765–70, 2003. doi:10.1073/pnas.2235465100.

Miguel G. Matias, Dominique Gravel, François Guilhaumon, Philippe Desjardins-Proulx, Michel Loreau, Tamara Münkemüller, and Nicolas Mouquet. Estimates of species extinctions from species-area relationships strongly depend on ecological context. Ecography, 37(5):431–442, 2014. doi:10.1111/j.1600-0587.2013.00448.x.

Nicolas Mouquet and Michel Loreau. Community patterns in source-sink metacommunities. The American Naturalist, 162(5):544–557, 2003. doi:10.1086/378857.

Otso Ovaskainen. Long-term persistence of species and the sloss problem. Journal of Theoretical Biology, 218(4):419–433, 2002. doi:10.1006/jtbi.2002.3089.

Otso Ovaskainen, Dmitri Finkelshtein, Oleksandr Kutoviy, Stephen Cornell, Benjamin Bolker, and Yuri Kondratiev. A general mathematical framework for the analysis of spatiotemporal point processes. Theoretical Ecology, 7(1):101–113, 2014. doi:10.1007/s12080-013-0202-8.

Henrique M. Pereira, Paul W. Leadley, V. Proenca, Rob Alkemade, Jörn P W Scharlemann, J. F. Fernandez-Manjarres, M. B. Araujo, Patricia Balvanera, Reinette Biggs, William W L Cheung, Louise Chini, H. David Cooper, Eric L. Gilman, S. Guenette, George C. Hurtt, Henry P. Huntington, Georgina M. Mace, Thierry Oberdorff, Carmen Revenga, Patrícia Rodrigues, Robert J Scholes, Ussif Rashid Sumaila, and Matt Walpole. Scenarios for global biodiversity in the 21st century. Science, 330(6010):1496–1501, 2010. doi:10.1126/science.1196624.

Pradeep Pillai, Michel Loreau, and Andrew Gonzalez. A patch-dynamic framework for food web metacommunities. Theoretical Ecology, 3(4):223–237, 2010. doi:10.1007/s12080-009-0065-1.

S. L. Pimm, C. N. Jenkins, R. Abell, T. M. Brooks, J. L. Gittleman, L. N. Joppa, P. H. Raven, C. M. Roberts, and J. O. Sexton. The biodiversity of species and their rates of extinction, distribution, and protection. Science, 344(6187):1246752–1246752, 2014. doi:10.1126/science.1246752.

Sona Prakash and André M. de Roos. Habitat destruction in mutualistic metacommunities. Theoretical Population Biology, 65(2):153–163, 2004. doi:10.1016/J.TPB.2003.10.004.

Michael Rosenzweig. Species diversity in space and time. Cambridge University Press, 1995.

Joel Rybicki and Ilkka Hanski. Species-area relationships and extinctions caused by habitat loss and fragmentation. Ecology Letters, 16(SUPPL.1):27–38, 2013. doi:10.1111/ele.12065.

Chris D. Thomas, Alison Cameron, Rhys E. Green, Michel Bakkenes, Linda J. Beaumont, Yvonne C. Collingham, Barend F. N. Erasmus, Marinez Ferreira de Siqueira, Alan Grainger, Lee Hannah, Lesley Hughes, Brian Huntley, Albert S. van Jaarsveld, Guy F. Midgley, Lera Miles, Miguel A. Ortega-Huerta, A. Townsend Peterson, Oliver L. Phillips, and Stephen E. Williams. Extinction risk from climate change. Nature, 427(6970):145–148, 2004. doi:10.1038/nature02121.

Patrick L. Thompson, Bronwyn Rayfield, and Andrew Gonzalez. Robustness of the spatial insurance effects of biodiversity to habitat loss. Evolutionary Ecology Research, 16:445–460, 2014.

Patrick L. Thompson, Bronwyn Rayfield, and Andrew Gonzalez. Loss of habitat and connectivity erodes species diversity, ecosystem functioning, and stability in metacommunity networks. Ecography, 40 (1):98–108, 2017. doi:10.1111/ecog.02558.

David Tilman, Robert M. May, Clarence L. Lehman, and Martin A. Nowak. Habitat destruction and the extinction debt. Nature, 371(6492):65–66, 1994. doi:10.1038/371065a0.

David Tilman, Clarence L. Lehman, and Chengjun Yin. Habitat destruction, dispersal, and deterministic extinction in competitive communities. Source: The American Naturalist The American Naturalist, 149(3):407–435, 1997.

Shaopeng Wang and Michel Loreau. Biodiversity and ecosystem stability across scales in metacommunities. Ecology Letters, 19(5):510–518, 2016. doi:10.1111/ele.12582.

Maxwell C. Wilson, Xiao-Yong Chen, Richard T. Corlett, Raphael K. Didham, Ping Ding, Robert D. Holt, Marcel Holyoak, Guang Hu, Alice C. Hughes, Lin Jiang, William F. Laurance, Jiajia Liu, Stuart L. Pimm, Scott K. Robinson, Sabrina E. Russo, Xingfeng Si, David S. Wilcove, Jianguo Wu, and Mingjian Yu. Habitat fragmentation and biodiversity conservation: key findings and future challenges. Landscape Ecology, 31(2):219–227, 2016. doi:10.1007/s10980-015-0312-3.

Kimberly A. With. Are landscapes more than the sum of their patches? Landscape Ecology, 31(5): 969–980, Jun 2016. doi:10.1007/s10980-015-0328-8.

Zhichao Xu, Yang Shen, and Jinbao Liao. Patch dynamics of various plant-animal interactions in fragmented landscapes. Ecological Modelling, 368:27–32, 2018. doi:10.1016/J.ECOLMODEL.2017.11. 017.

